# Kidney kallikrein-1 contributes to cleavage of gamma-ENaC *in vivo*

**DOI:** 10.1101/2025.09.19.677394

**Authors:** Joshua Curry, Xiao Tong Su, Qi Wu, Yujiro Maeoka, Chao-Ling Yang, Eric Delpire, Robert A. Fenton, Paul A. Welling, David H. Ellison

## Abstract

The epithelial sodium channel (ENaC) is essential for sodium reabsorption and potassium homeostasis in the distal nephron, where its activity is controlled by mineralocorticoid signaling and downstream proteolytic processing of channel subunits. While cleavage of the γ-ENaC subunit has been implicated in aldosterone-mediated sodium transport, the identity of mineralocorticoid receptor (MR)-regulated proteases responsible for this process remains uncertain. Here, we investigated the role of kallikrein-1 (encoded by *Klk1*), a serine protease expressed in the connecting tubule and cortical collecting duct (CNT/CCD), as a mediator of ENaC activation. Using CRISPR/Cas9, we generated a conditional *Klk1*-floxed allele and established mice with CNT/CCD-specific deletion of *Klk1* by crossing with *Calb1*-Cre (CNT-*Klk1*^-/-^). On a low sodium, high potassium diet, CNT-*Klk1*^-/-^ mice exhibited ∼85% less renal kallikrein-1 expression, yet maintained normal serum electrolytes, urinary potassium excretion, and aldosterone responses. Western blot analysis revealed significantly less cleavage of γ-ENaC and α-ENaC in CNT-*Klk1*^-/-^ kidneys, accompanied by more total NCC abundance. Despite impaired ENaC proteolysis, amiloride-sensitive sodium excretion was preserved, indicating intact ENaC function. These findings identify renal kallikrein-1 as a protease that contributes to ENaC subunit processing *in vivo*. However, the absence of overt sodium or potassium handling defects in CNT-*Klk1*^-/-^ mice suggests that kallikrein-1 deficiency is not sufficient to disrupt overall ENaC function, likely due to compensatory mechanisms from redundant proteolytic or non-proteolytic pathways. Together, our results refine the role of kallikrein-1 as a modulator, rather than a sole determinant, of ENaC activation and highlight the complexity of aldosterone-dependent sodium transport in the distal nephron.

**New & Noteworthy:** Using a novel connecting tubule / cortical collecting duct specific kallikrein-1 knockout model, we show that γ- and α-ENaC cleavage is impaired by loss of renal kallikrein-1 without major disturbances in sodium or potassium handling. These findings highlight redundancy among ENaC regulatory pathways and suggest that proteolytic cleavage, while biochemically evident, may not be an accurate marker of ENaC-mediated sodium transport under physiological stress.

## Introduction

Sodium and potassium homeostasis is regulated in part by the epithelial sodium channel (ENaC), which mediates apical sodium reabsorption in the distal nephron. ENaC activity is tightly controlled through multiple mechanisms, including transcriptional regulation, trafficking to the plasma membrane, and modulation of channel open probability (1). *In vivo*, ENaC activity is increased by both dietary sodium depletion and activation of the mineralocorticoid receptor (MR) by aldosterone (2, 3). The MR regulates ENaC activity in the kidney by multiple mechanisms including by modulating ENaC subunit transcription and channel trafficking, and by controlling proteolytic cleavage of ENaC subunits (4-6). Substantial evidence links proteolytic cleavage of ENaC to increased channel activity *in vitro* (7, 8), and studies using animal models suggest that cleavage of ENaC is mediated by aldosterone/MR (5, 9, 10). Despite the known association between MR signaling and γ-ENaC cleavage in the distal nephron, the identity of the MR-regulated protease(s) responsible for this cleavage event remains uncertain.

ENaC is composed of α, β, and γ subunits, which form a 1:1:1 heterotrimeric channel (11). Both α and γ subunits undergo proteolytic cleavage at sites flanking a gating release of inhibition by proteolysis (GRIP) domain. Proteolytic cleavage at these sites releases the inhibitory GRIP peptide and increases channel open probability *in vitro* (11-13). Intriguingly, the sites of cleavage appear to have independently co-evolved for each isoform during the vertebrate transition to terrestrial life (14). Cleavage of both α-ENaC sites and the proximal site of γ-ENaC is mediated by furin cleavage at consensus sequences during intracellular trafficking through the trans-Golgi network (7). A subset of ENaC channels resides within the plasma membrane with low open probability-these channels require extracellular proteases to cleave γ-ENaC at its distal site and release the γ-ENaC inhibitory peptide to increase open probability (15, 16). This distal site is recognized by several serine proteases, including prostasin, originally identified as channel-activating protein 1, and kallikrein-1 (13, 17).

Early studies suggested that prostasin (then called channel activating protein-1) was responsible for ENaC cleavage (18, 19). Importantly, prostasin expression appears to be MR-independent, as MR deletion did not alter prostasin levels in mice (5). In contrast, kallikrein-1—a serine protease expressed at high levels in the aldosterone sensitive distal nephron—has been shown to be MR-responsive. In human subjects, for instance, urinary kallikrein activity increased with fludrocortisone (an MR agonist) and decreased with spironolactone (an MR antagonist), suggesting regulation by MR (20). Consistent with this, we recently identified *Klk1*, the gene for kallikrein-1, as a differentially regulated transcript in the distal nephron of mice with tubule-specific deletion of MR (manuscript in preparation).

The kallikrein–kinin system (KKS) is a paracrine pathway in which tissue and plasma kallikreins hydrolyze kininogens to locally generate kinins, leading to cell signaling responses that regulate vascular tone, renal blood flow, and sodium handling (21). Within the kidney, kallikrein-1 is the predominant tissue kallikrein isoform, primarily synthesized in the connecting tubule (CNT) and secreted into the urine (22). *In vitro*, urinary kallikrein exposure induces concurrent amiloride-sensitive currents and cleavage of γ-ENaC in *Xenopus* oocytes expressing wild type ENaC, but not in those with a mutated distal cleavage site in the γ subunit (17). A global *Klk1* knockout mouse model was previously generated that provided important insights into kallikrein-1 biology. These animals exhibited abnormalities in vascular and cardiac function (23, 24), as well as defective γ-ENaC cleavage and post-prandial hyperkalemia (3, 25). Despite these biochemical and systemic changes, global Klk1 deficiency did not produce a baseline defect in renal sodium handling, perhaps because of confounding effects of systemic kallikrein–kinin deficiency on vascular tone and bradykinin signaling. The present work therefore isolates renal kallikrein-1 function to define its role in MR-dependent distal transport regulation, including γ-ENaC cleavage, independent of systemic KKS actions.

## Materials and Methods

### Generation of Kallikrein-1-flox mice

All animal studies were approved by the Oregon Health and Science University (Protocol TR02_IP00000286) and Vanderbilt University Medical Center Animal Care and Use Committee. A conditional kallikrein-1 (*Klk1*) mutant mouse was generated using CRISPR/Cas9 gene editing to target the *Klk1* gene located on mouse chromosome 7. Exon 2 was selected as the target site, with guide RNAs designed within the first (AATAGTACACCTCCCTATCT) and second (CTGTACATAAACCTGGATGG) introns. Guide RNAs, recombinant Cas9, and a 656-base single-stranded repair oligonucleotide were microinjected into 300 x 0.5-day C57BL/6J embryos. Surviving embryos were implanted into 11 pseudo-pregnant females, resulting in the birth of 33 pups. Genomic DNA extracted from tail biopsies was sequenced to assess CRISPR-mediated modifications. The following genotypic distribution was observed: 13 wild-type alleles, 2 with cut-site deletions, 6 with indels, 2 mosaic mice, 4 containing a single loxP site, and 5 pups carrying both loxP sites (#6, #8, #9, #10, and #12). Sequence verification of several hundred base pairs around each loxP site was performed using primers flanking the recombination arms to ensure proper locus integration. Line #8 was selected and backcrossed to C57BL/6J for three additional generations to eliminate potential off-target effects before generating homozygous mice.

### Generation of kidney-specific Kallikrein-1 knockout models

We recent reported a Calbindin 1(*Calb1*)-Cre recombinase expressing model in a study utilizing a nuclear reporter line to isolate and sequence all nuclei expressing *Calb1* transcripts. While this study was designed to isolate CNT and cortical collecting duct cell populations, our results suggest that *Calb1* is expressed during development all along the distal nephron, including the distal convoluted tubule, CNT, and collecting duct, but not the thick ascending limb (*in submission*). Thus, for conditional deletion of kallikrein-1 in the distal nephron, *Klk1* flox mice were crossed with *Calb1*-Cre mice to generate animals with distal tubule-specific knockout of kallikrein-1. Experimental animals used in this study include constitutive *Klk1*^flox/flox^ *Calb1*-Cre mice (hereby referred to as CNT *Klk1*^-/-^) and *Klk1*^*flox*/flox^ *Calb1*-Cre negative littermate controls (hereby referred to as CNT *Klk1*^+/+^). All mice used in experiments were between 8-17 weeks of age. Genotyping was performed by quantitative PCR by Transnetyx (Memphis, TN).

### Metabolic cage studies

All animals were housed in a temperature-controlled room under standard light/dark cycle (12:12 hours) with free access to food and drinking water, unless otherwise specified. Animals were fed a gel diet (AIN-93G Rodent Diet with 0.03% Na^+^ and 0.36% K^+^; Animal Specialties Incorporated, 1813804-103) with NaCl and KCl supplementation as specified. Control diet was prepared with 0.22% Na^+^ and 0.9% K^+^ (control diet) and low sodium, high K^+^ diet with 0.03% Na^+^, 3% K^+^ (Low Na/Hi K). Animals were housed in metabolic cages and urine was collected under water–saturated light mineral oil for 24 hours.

### Amiloride challenge

CNT *Klk1*^-/-^ and CNT *Klk1*^+/+^ mice were placed on the Low Na/HiK diet for 5 days during the amiloride response test. On Day 4, animals were injected intraperitoneally with vehicle (0.9% saline) and then placed in metabolic cages for a 6-hour fasting urine collection, while maintaining free access to water. Following collection, animals were returned to standard cages with continued access to the experimental diet. On Day 5, the same animals received intraperitoneal injection of amiloride hydrochloride (40 µg per 25 g body weight), followed by another 6-hour urine collection under identical conditions to the vehicle collection.

### Urine and blood analysis

Whole blood was collected under anesthesia (isoflurane) via cardiac puncture, and chemistry was measured using an i-STAT point-of-care autoanalyzer (Chem 8+, Abbott). Urine Na^+^ and K^+^ contents were determined by flame photometry. Urine calcium content was determined by colorimetric assay (Pointe Scientific, C7503-480). Urine aldosterone was measured using competitive enzyme-linked immunosorbent assay (ELISA, IBL America #IB79134).

### Western blot

Following cardiac puncture, kidneys were harvested and snap frozen in liquid nitrogen to be stored at - 80°C until homogenization. Kidneys were homogenized using a Potter-Elvehjem homogenizer in 1 mL ice-cold homogenization buffer ((300 mM sucrose, 50 mM Tris·HCl, pH 7.4, 1 mM EDTA, 1 mM EGTA, 1 mM NaVO4, 50 mM NaF, 1 mM dithiothreitol, 1 mM phenylmethylsulfonyl fluoride, 1 mg/ml aprotinin and 4 mg/ml leupeptin) with added PhosSTOP phosphatase inhibitor cocktail (Roche, Cat. No. 04906837001). Homogenate was centrifuged, and supernatant was transferred to a new tube. After total protein quantification, 20-40 µg protein per sample was separated on 7.5% Criterion TGX Stain-Free protein gels (Bio-Rad, Hercules, CA). Membranes were then washed, incubated with an HRP–coupled secondary antibody (1:5000; Invitrogen, Carlsbad, CA), washed again, and imaged using a Western Lightning Plus Kit (Revivity Health Sciences, Waltham, MA). Antibodies and dilutions for anti-kallikrein-1, γ-ENaC (26), α-ENaC (27), calbindin (28), total (29) and phosphorylated (30) NCC are listed in Supplementary Table 1.

### Immunofluorescence of paraffin embedded kidney tissue

Kidneys were harvested following exsanguination and incubated in 10% neutral buffer formalin overnight at 4°C prior to storage in 70% ethanol. Tissue was processed in paraffin blocks by the Oregon Health and Science University Histopathology Core Facility and sectioned onto slides (5-µm sections). Paraffin was removed by immersion for 5 min in 100% xylene and then kidney tissue was rehydrated with successive immersion in 100%, 90%, and 70% ethanol solutions. Antigen retrieval was performed by microwaving tissue in a citrate-based antigen unmasking solution (Vector Laboratories, Cat. No. H-3300). Sections were blocked with 1% BSA in PBS for 30 min followed by an incubation with primary antibody diluted in blocking buffer overnight at 4°C. Sections were incubated with fluorescent dye-conjugated secondary antibody (1:1000; Invitrogen, Carlsbad, CA) diluted in blocking buffer for 1 hour at room temperature before being mounted with Prolong Diamond Antifade Mountant with DAPI (Invitrogen, Carlsbad, CA). Antibodies and dilutions used are listed in Supplementary Table 1.

### Urine proteomics

For analysis of urinary protein excretion, a group of CNT-*Klk1*^-/-^ and CNT Klk1^+/+^ littermates (n=4 males of each genotype) were placed on the Low Na/Hi K diet for 5 days. 24-hour urine was collected on day 5 under water-saturated mineral oil with 10x protease inhibitors (cOmplete™ mini, Sigma) and phosphatase inhibitors (PhosSTOP™, Sigma) dissolved in pure water added to the collection tube. Each urine sample was acetone-precipitated and dissolved in 200 µL of 1% SDS in 1x PBS with protease (cOmplete™ mini, Sigma) and phosphatase (PhosSTOP™, Sigma) inhibitors, then sonicated thoroughly by a probe sonicator. The sample was then centrifuged at 16,000g for 30 minutes and the supernatant kept for filter-aided sample preparation (FASP). Specifically, supernatants from each sample were loaded to a Vivacon-30 kDa spin column (Sartorius) and centrifuged at 16,000 g to remove all liquids. The columns were then washed three times with a UA buffer containing 8 M urea and 100 mM triethylammonium bicarbonate (Thermo Fisher). Subsequently, proteins were reduced with 50 mM dithiothreitol (Thermo Fisher) for 1 hour at 37□J°C, followed by alkylation with 50 mM iodoacetamide (Thermo Fisher) for 20 minutes at room temperature in the dark. After each step, centrifugation was performed to remove all liquids. Another UA buffer wash was carried out before adding 5 µl of Lys-C (Thermo Fisher) solution at an enzyme-to-protein ratio of 1:50, dissolved in UA buffer. The mixture was incubated at 37□J°C for 2 hours, after which 50 µl of trypsin (Thermo Fisher) solution (enzyme-to-protein ratio of 1:25, dissolved in 100 mM triethylammonium bicarbonate) was added. Digestion proceeded overnight at 37□J°C. Next morning, 4 µL of 10% TFA (Thermo Fisher) was added to the samples followed by desalting using C18 desalting tips (Thermo Fisher). After desalting, samples were lyophilized in a SpeedVac until fully dried.

### LC-MS analysis

Samples were analyzed by a Vanquish Neo HPLC coupled with an Orbitrap Ascend mass spectrometer (MS) and a FAIMS Pro Duo interface (Thermo Fisher). A 30-minute gradient with an active separation window of 24.5 minutes of 5-40% of B (80% ACN and 0.1% FA) was used to separate the peptides on a 50-cm long µPAC column (Thermo Fisher). MS analysis was done in Data Independent Acquisition (DIA) mode, with FAIMS CV of -50. Parameters for MS1 scans include scan range of 380-980, Orbitrap resolution of 120K, maximum ion injection time of 251ms, normalized AGC of 250%. Parameters for MS2 scans include DIA window size of 40 Da, Orbitrap resolution of 120K, maximum ion injection time of 251ms, normalized AGC of 1000%, collision energy of 30%, number of scan events = 15, loop count of 5 (one MS1 scan inserted between every 5 MS2 scans), and loop time of 3 seconds.

### Data analysis

The MS raw data was analyzed by the direct DIA+ mode in Spectronaut (v19, Biognosys) against a mouse uniport database downloaded on Jan 6, 2025. Peptide and protein false discovery rates were set at 1%. All parameters were default except the quantification level was set to MS1. Identified proteins and the quantification matrix were exported to a csv file for downstream data analysis. For proteomics analysis, significance was defined as p<0.05 (uncorrected). Although false discovery rate (FDR) adjustment was performed, no proteins remained significant at q<0.05 with n=4 per group.

### Statistical Analyses

Null-hypothesis testing was performed by unpaired *t* test, two-way ANOVA, or two-way ANOVA for repeated measures as indicated. For Table 1 and Supplementary Table 2, *p*-values were adjusted for multiple comparisons using the Benjamini-Hochberg (FDR) method. For the amiloride challenge test, urine sodium to potassium ratios were log-transformed to normalize data. The response to amiloride was quantified as the difference between log-transformed values from amiloride and vehicle urine collections (Δlog(UNa/UK). Fold changes were then calculated by back-transforming these data using the equation: Fold Change = 10^Δlog(UNa/UK)^.

**Table 1.** Blood parameters of Male Mice on Control and Low Sodium, High Potassium Diet.

| Males | Control |  |  | CNT *Klk1*<sup>-/-</sup> |  |  | P |
| --- | --- | --- | --- | --- | --- | --- | --- |
|  | Mean | SEM | *n* | Mean | SEM | *n* | Value |
| **Parameter** |  |  |  |  |  |  |  |
| ***Control Diet*** |  |  |  |  |  |  |  |
| Na | 142.1 | 0.93 | 8 | 141.8 | 0.86 | 5 | 0.83 |
| K | 3.71 | 0.13 | 8 | 3.86 | 0.19 | 5 | 0.51 |
| Cl | 112.6 | 1.64 | 8 | 112.4 | 1.63 | 5 | 0.94 |
| TCO<sub>2</sub> | 18.4 | 0.56 | 8 | 18.0 | 0.63 | 5 | 0.65 |
| iCa | 1.26 | 0.02 | 8 | 1.26 | 0.02 | 5 | 0.99 |
| Glu | 289.6 | 20.9 | 8 | 313.6 | 32.8 | 5 | 0.53 |
| BUN | 20.1 | 0.97 | 8 | 19.0 | 1.14 | 5 | 0.49 |
| Hct | 37.3 | 0.73 | 8 | 38.4 | 1.50 | 5 | 0.47 |
| ***Low Na/High K Diet*** |  |  |  |  |  |  |  |
| Na | 144.7 | 1.74 | 7 | 139.7 | 2.22 | 7 | 0.10 |
| K | 4.34 | 0.20 | 7 | 4.73 | 0.23 | 7 | 0.22 |
| Cl | 115.4 | 2.69 | 7 | 112.0 | 2.19 | 7 | 0.35 |
| TCO<sub>2</sub> | 19.3 | 0.71 | 7 | 18.8 | 0.84 | 7 | 0.66 |
| iCa | 1.28 | 0.01 | 7 | 1.22 | 0.03 | 7 | 0.08 |
| Glu | 257.7 | 25.8 | 7 | 302.4 | 16.9 | 7 | 0.17 |
| BUN | 21.1 | 1.26 | 7 | 22.9 | 2.60 | 7 | 0.54 |
| Hct | 37.3 | 1.06 | 7 | 38.9 | 1.16 | 7 | 0.33 |

## Results

### Generation and Validation of *Klk1* Flox Mice

To investigate the functional role of renal kallikrein-1 in ENaC activation and electrolyte handling *in vivo*, we generated a conditional *Klk1* knockout mouse model to delete Klk1 in the CNT, the major site of kallikrein-1 synthesis in the renal tubule (22). Using CRISPR/Cas9 gene-editing technology, we inserted *loxP* sites flanking exon 2 of the Klk1 gene (Figure 1A). PCR-based genotyping confirmed successful insertion of the loxP sites (Figure 1B). *Klk1* Flox mice were then bred with mice expressing Cre recombinase under the control of the calbindin-1 (*Calb1*) promoter (CB-Cre), enabling CNT- and CCD-specific knockout of *Klk1* (CNT-*Klk1*^-/-^). Immunofluorescent imaging of kidney sections demonstrates qualitative loss of kallikrein-1 protein expression in calbindin-positive tubules in the distal nephron of CNT-*Klk1*^-/-^ mice, with scant positive staining in other tubule segments (Figure 1C).

**Figure 1.**
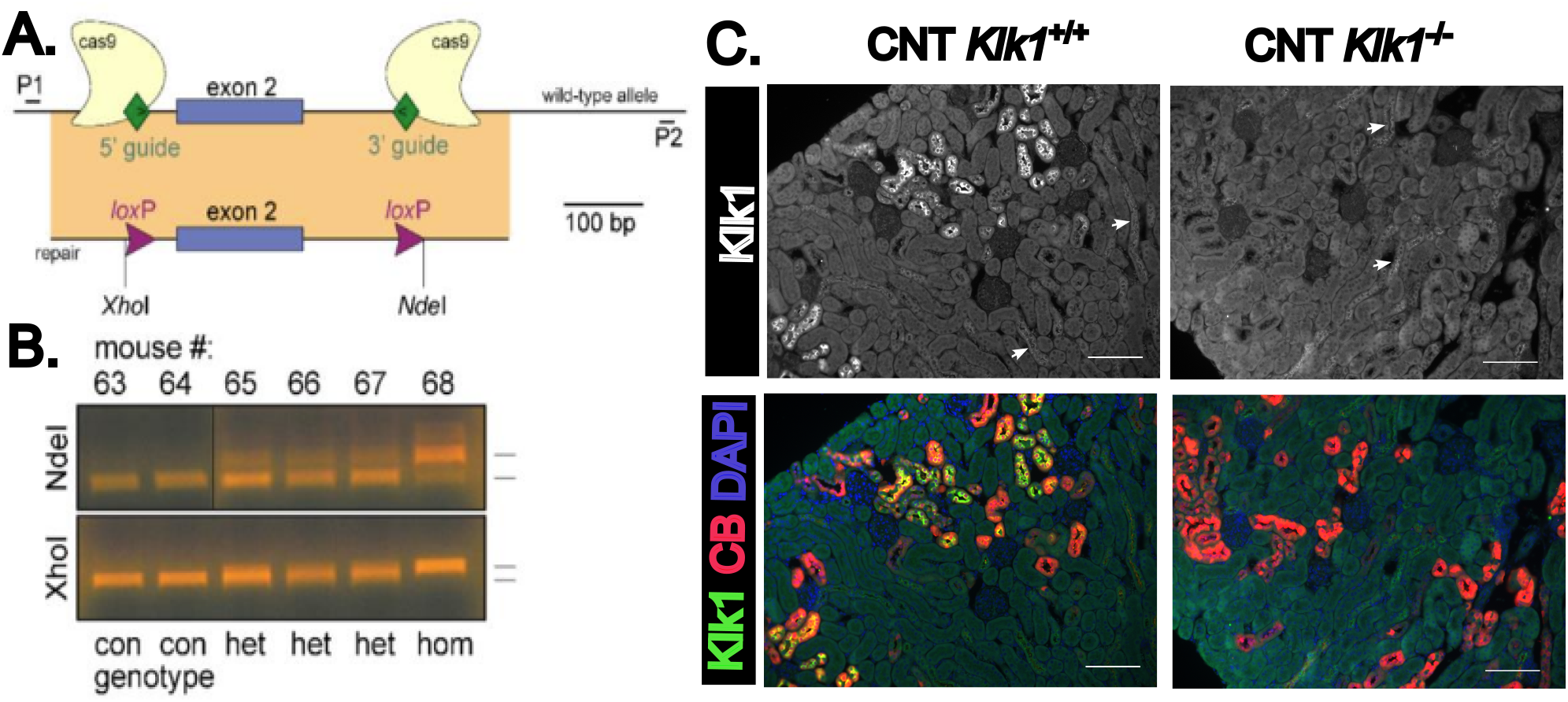
Generation of Klk1-flox mouse. A. Guide RNAs were selected upstream and downstream of exon 2 to target cas9 to the *Klk1* locus. Repair DNA introduces loxP sites as well as upstream XhoI and downstream NdeI restriction enzyme sites. B. WT and mutant alleles amplified using P1 and P2 primers and the PCR fragments digested with either NdeI (top) or XhoI (bottom). Two wild type (con), three heterozygotes (het) and one homozygote (hom) mouse were identified in a litter of six pups. The alleles from the homozygouse mouse were sequenced to confirm loxP sites. C. Connecting tubule specific *Klk1* deletion was confirmed using immunofluorescence of paraffin-embedded kidney tissue, which demonstrated strong kallikrein-1 staining of calbindin-positive tubules (CNT *Klk1*^+/+^) and loss of kallikrein-1 staining in CNT *Klk1*^-/-^ mice. Very faint kallikrein-1 staining was detected in calbindin-negative tubule segments (arrows). Scale bar = 100_u_m.

### Mice with Deletion of Kallikrein-1 in the Aldosterone Sensitive Distal Nephron Maintain Potassium Excretion Despite Loss of Renal Kallikrein-1

To assess the physiological impact of CNT/CCD-specific *Klk1* deletion, we fed CNT-*Klk1*^-/-^ mice and CNT Klk1^+/+^ littermates 5 days of normal (control) or low sodium, high potassium (low Na/hi K) diet. There were no differences detected in food intake or urine volume (Table S2). CNT-*Klk1*^-/-^ mice exhibited ∼85% or less kallikrein-1 protein expression regardless of diet, as determined by densitometry of Western blots (Figure 2A-B, males; S1A-B, females). As expected, 24-hour urine aldosterone excretion was elevated in the low Na/hi K groups compared with control diet (Figure 2C). There was no difference in serum potassium levels between knockout and control animals (two-way ANOVA, genotype p=0.164) on either diet (Figure 2D, males; S1D, females). There were no other notable differences in blood parameters in either males (Table 1) or females (Table S3). Similarly, urine sodium, urine potassium, urine volume, and urine calcium did not differ between genotypes (Figure 2E-H, males; S1E-H, females). These results suggest that CNT/CCD kallikrein-1 is not essential for potassium excretion even under conditions of dietary stress to stimulate aldosterone secretion.

**Figure 2.**
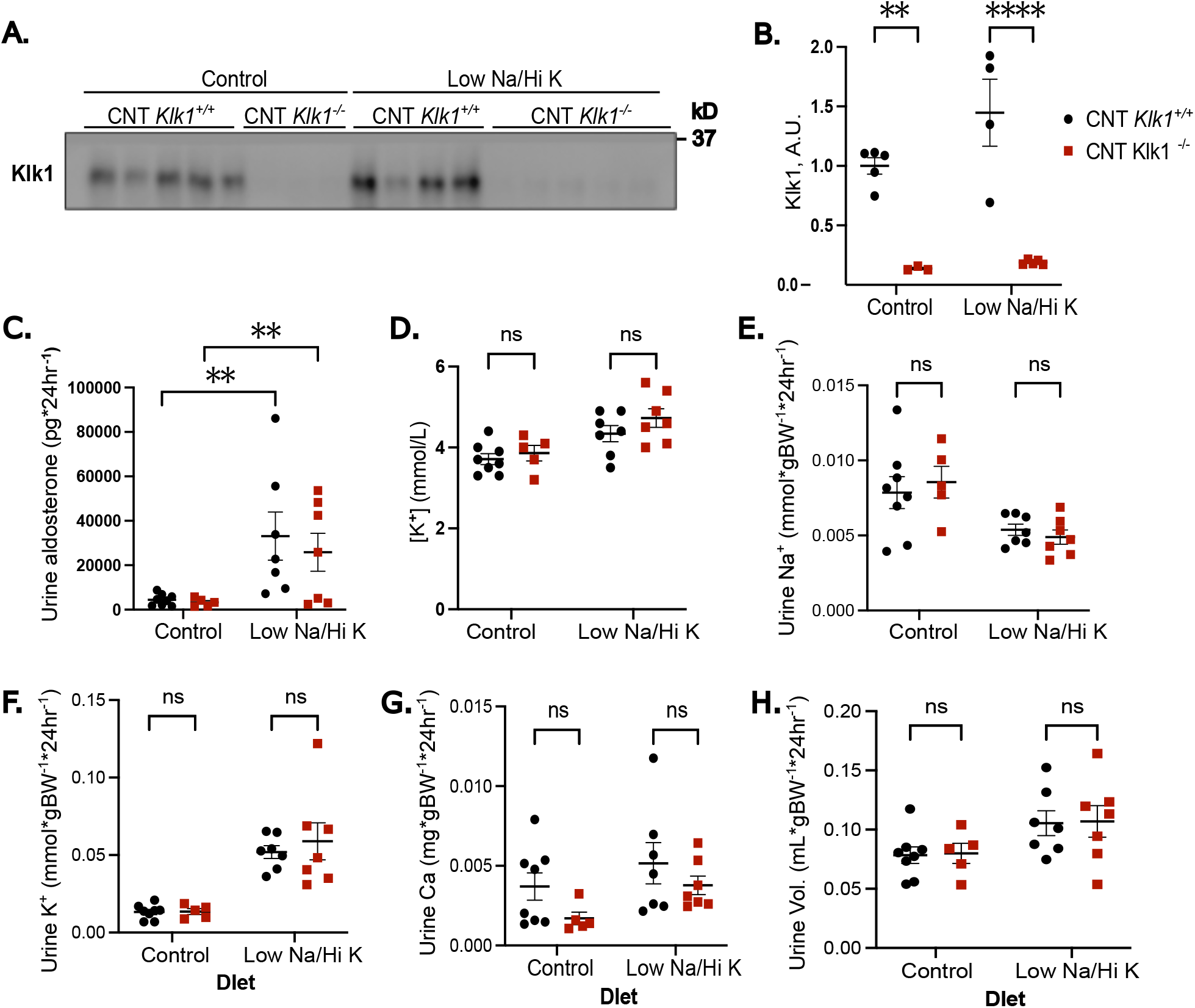
Mice with Distal Nephron-Specific Deletion of Kallikrein-1 Maximally Conserve Potassium in response to a Low Sodium, High Potassium Diet. Males, A. Western blot and B. quantification of kallikrein-1 abundance demonstrating a significant reduction in kallikrein-1 expression in the *Klk1*^flox/flox^ *Calb1*-Cre mice (CNT *Klk1*^-/-^) follow 5 days on a normal salt diet (Control) or low sodium, high potassium diet (Low Na/Hi K). On the final day of dietary challenge, mice were placed in metabolic cages for 24-hour urine collection prior to blood and kidney tissue collection. C. 24-hour urinary aldosterone excretion demonstrates increases aldosterone in response to Low Na/Hi K. D. Serum potassium did not differ in CNT *Klk1*^-/-^. Likewise, we were unable to detect a difference in urinary sodium (E), potassium (F), calcium (G), or urine volume (H) under either dietary condition. Results were analyzed by two-way ANOVA with Boferroni post-hoc correction for multiple comparisons. ^*^\**P*<0.01; ^***^\**P*<0.0001.

### CNT-*Klk1*^-/-^ Mice Exhibit Defective ENaC Cleavage in Response to a Low Sodium, High Potassium Diet

To determine the role of kallikrein-1 in ENaC activation, we performed Western blot analysis on kidneys from CNT-*Klk1*^-/-^ and control mice subjected to the short-term low Na/hi K diet. Our results revealed that CNT-*Klk1*^-/-^ exhibited significantly less cleaved and trends towards less uncleaved (p=0.0681) γ-ENaC compared with controls on the low Na/hi K diet (two=way ANOVA; Figure 3A-C, males; S2A-C, females). After applying Bonferroni correction for multiple comparisons, the adjusted p-value for cleaved γ-ENaC remained significant in both males (p = 0.0015) and females (p = 0.0179) for a genotype difference on the low Na/Hi K diet (Figure 3A-C, males; S2A-C, females). Unexpectedly, in addition to the predicted reduction in cleaved γ-ENaC, we observed significantly decreased α-ENaC cleavage in males (two-way ANOVA; Bonferroni correction for multiple comparisons p=0.0038) (Figure 3A, D-E, males; S2A, D-E, females). In addition to ENaC, total (tNCC) protein was higher in males (two-way ANOVA with Bonferroni correction, p = 0.0030) and phosphorylated NCC (pNCC) trended higher (p = 0.072) (Figure 3A, F-G). The ratio of tNCC to pNCC remained unchanged in CNT-*Klk1*^-/-^ males (low Na/hi K diet, CNT Klk1^+/+^ 0.926+/-0.035 vs. CNT-*Klk1*^-/-^ 0.861+/-0.030). In females, no differences in tNCC or pNCC were detected (Figure S2A, F-G).

**Figure 3.**
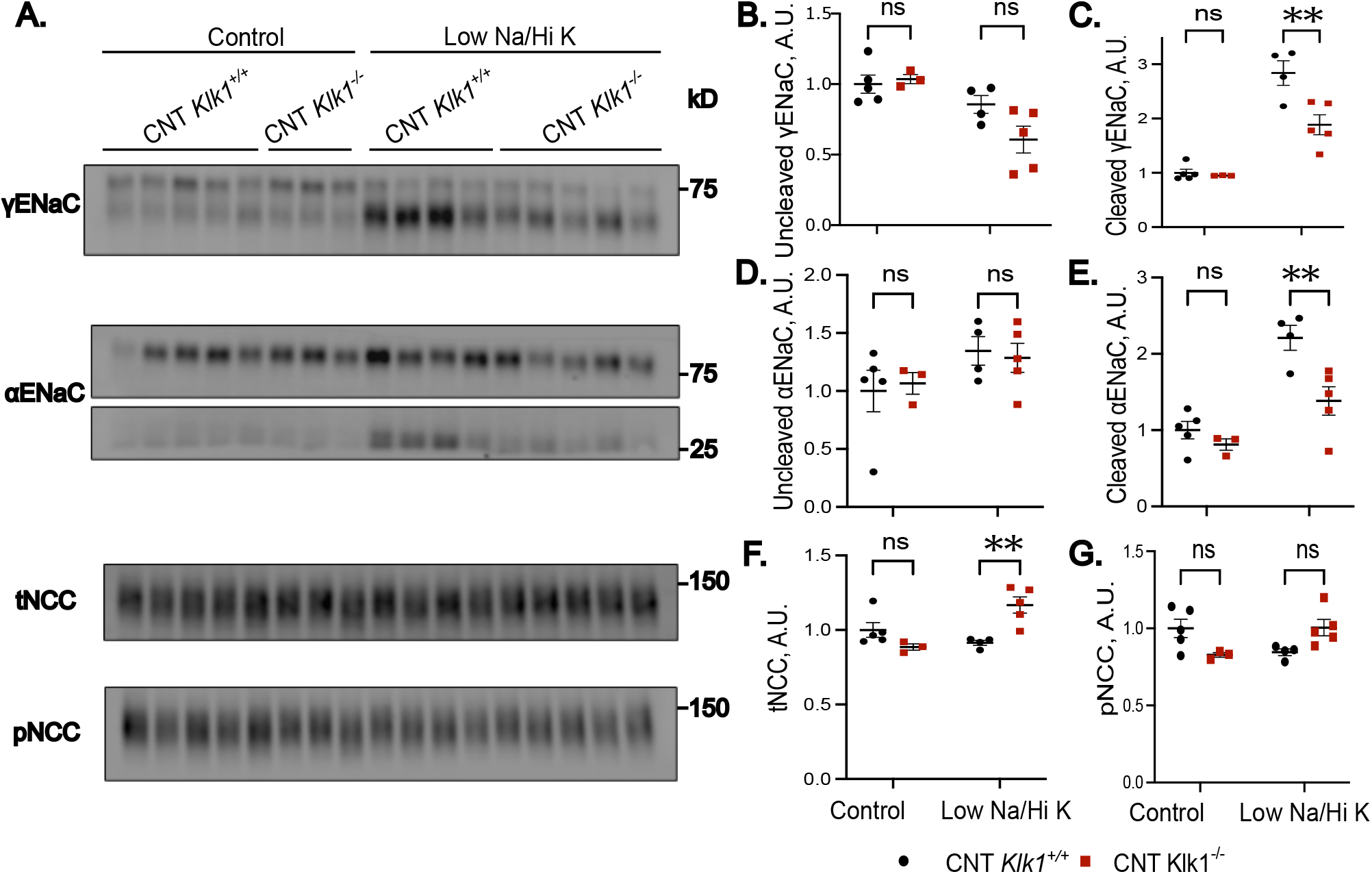
Loss of Distal Kallikrein-1 Leads to Reduced Cleavage of ENaC. A. Western blot of γ-ENaC, α-ENaC, total NCC (tNCC) and phosphorylated NCC (pNCC) abundance. B-G. Quantification of western blot results. A significant reduction in cleaved γ-ENaC (C), α-ENaC (E), and tNCC (F) was detected in CNT-*Klk1*^-/-^ mice following challenge with a low sodium, high potassium diet (Low Na/Hi K). Results were analyzed by two-way ANOVA followed by Bonferroni multiple comparison correction. ^*^\**P*<0.01.

### Despite Defective ENaC Cleavage, CNT-*Klk1*^-/-^ Mice Retain an Intact Amiloride Response

To test the functional difference in ENaC reabsorption following the low Na/hi K dietary challenge, we performed an amiloride response test. Amiloride resulted in a large increase in urine sodium excretion compared with vehicle (two-way repeated measures ANOVA, Figure 4A, males; Figure 4B, females). Despite impaired γ-ENaC cleavage, CNT-*Klk1*^-/-^ mice did not exhibit differences in amiloride responsiveness compared with control mice, as assessed by fold change of urine sodium to potassium ratio (Figure 4C, males; 4D, females).

**Figure 4.**
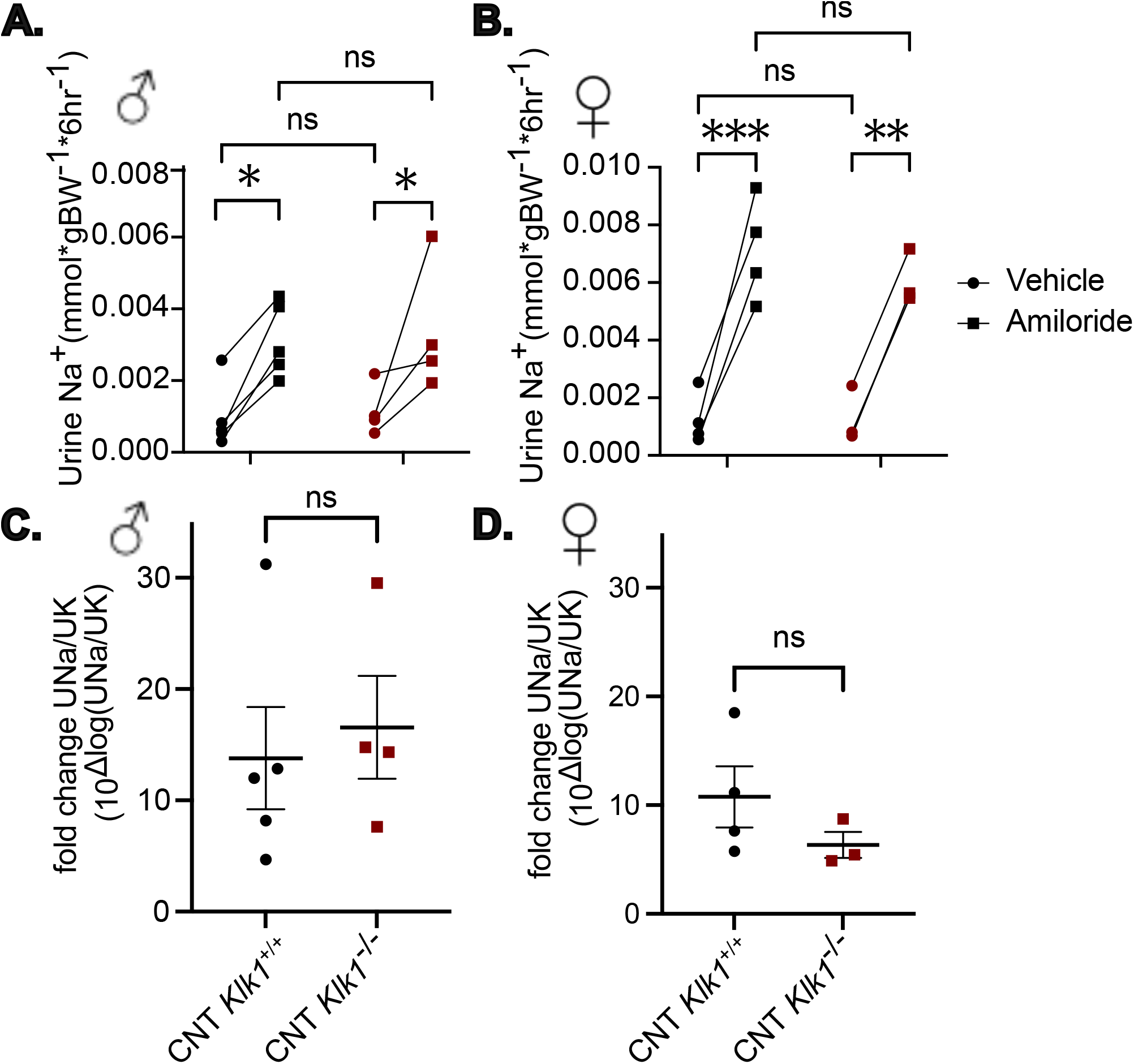
Despite defective ENaC Cleavage, Amiloride Response is Preserved in CNT-*Klk1*^-/-^. The determine the functional impact of impaired ENaC cleavage in CNT *Klk1*^*-/-*^, amiloride response test was performed. Animals were fed a low sodium, high potassium diet for 4 days prior to injection with vehicle (0.9% NaCl) followed by 6-hour urine collection. Animals were then returned to standard cages with access to the experimental diet. The following day, animals were injected with amiloride (40 µg 25 g^−1^ body weight in vehicle) followed by another 6-hour urine collection. Sodium excretion in males (A) and females (B) increased following challenge with amiloride, compared with vehicle, but no difference in urinary excretion between genotypes was detected by 2-way ANOVA with repeated measures. The urinary sodium to potassium ratio was analyzed following log-transform to normalize data. Fold change increase in urine sodium to potassium ratios in response to amiloride are shown for males (C) and females (D). There was no difference in amiloride response between genotypes. These results were analyzed by two-tailed *t-*test. \**P*<0.05; ^*^\**P*<0.01; ^**^\**P*<0.001.

### Urine proteomics did not identify compensatory increases in urine proteases in CNT-*Klk1*^-/-^

Given the normal amiloride response in CNT-*Klk1*^-/-^ mice, we hypothesized that loss of kallikrein-1 might be offset by increased secretion of alternative urinary proteases. To test this, we performed quantitative urine proteomics on male mice following 5 days of low Na/hi K diet (Figure 5A). As expected, urinary kallikrein-1 was markedly reduced in CNT-*Klk1*^-/-^ mice (∼86%; Figure 5B). However, a large amount of kallikrein-1 protein was detected in the urine, relative to other detected proteases. No compensatory increase was detected in other urinary proteases, including the murine kallikrein-1 analogue Klk1b5 or prostasin (Prss8; Figure 5B). We were unable to detect urinary Tmprss2 or Tmprss4 in this study.

**Figure 5.**
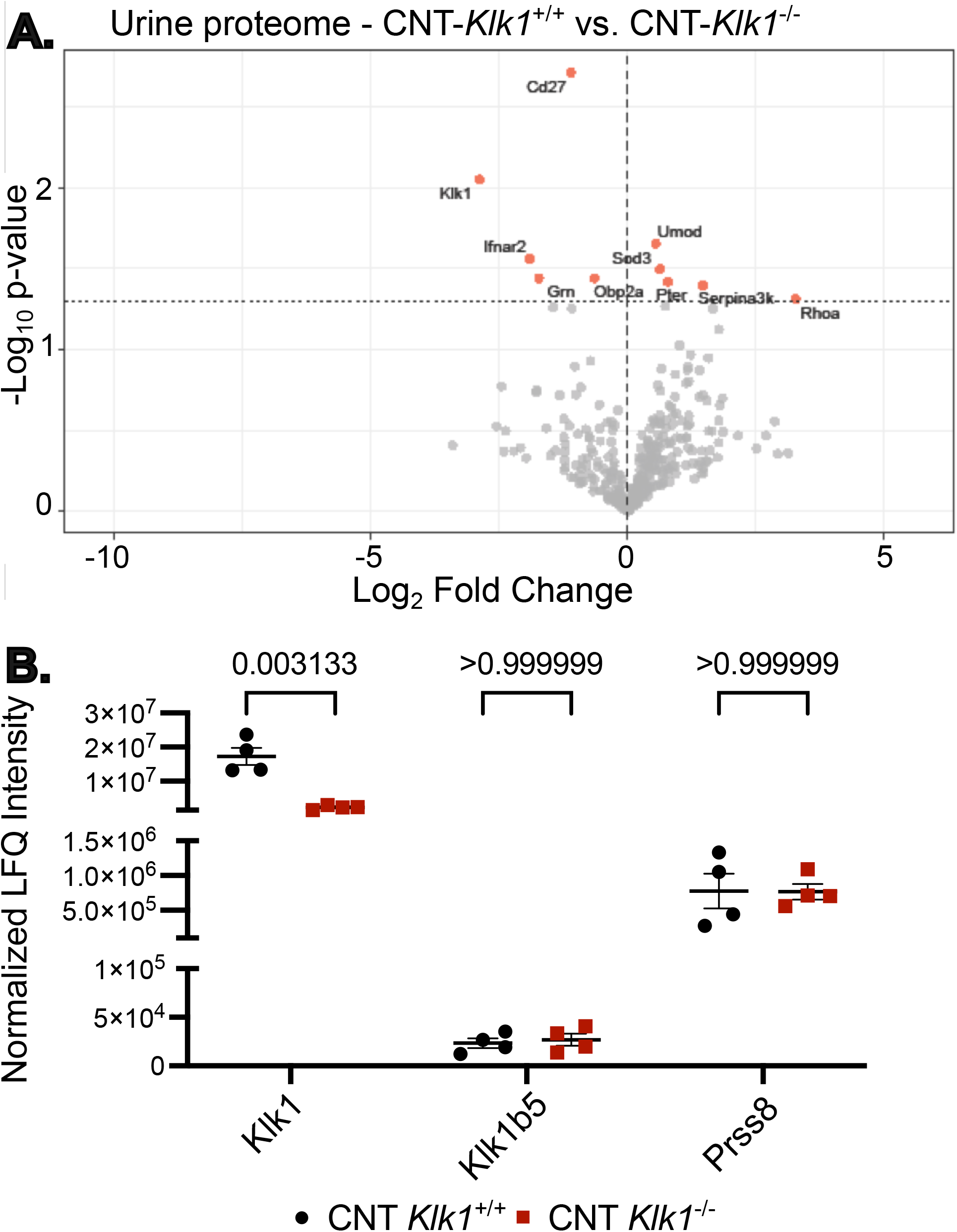
Urinary proteomics confirms reduction in urinary kallikrein-1 protein excretion in CNT *Klk1*^-/-^ without compensatory increases in prostasin or kallikrein isoforms. (A) Volcano plot of label-free quantification (LFQ) proteomics comparing urinary protein abundance in CNT-*Klk1*^-/-^ versus CNT *Klk1*^+/+^ mice. Each point represents a protein group, plotted as log_2_ fold change against -log_10_ p-value from an unpaired *t*-test. Horizontal dotted line indicates p = 0.05, and vertical dashed line indicates no change. Red points denote significantly altered proteins (p<0.05). Selected proteins are labeled. (B) Individual normalized LFQ intensity plots for selected proteins, analyzed by two-tailed *t*-test. Each point represents one mouse; error bars indicate mean +/-SEM. Klk1 is shown, confirming loss of urinary kallikrein-1 excretion in CNT-*Klk1*^-/-^. Klk1b5, another kallikrein family member, is not different, nor is Prss6 (prostasin). Numerical *P*-values are shown.

## Discussion

In this study, we identify kidney tubule-produced kallikrein-1 as a candidate protease that contributes to the proteolysis of ENaC channels in the distal nephron in response to mineralocorticoid receptor activity. Combined with our recent transcriptomic data (in submission), we demonstrate that kidney kallikrein-1 expression is regulated by MR and contributes to the post-translational processing of ENaC subunits *in vivo*. These findings suggest that kallikrein-1 acts as a downstream effector in MR-mediated sodium reabsorption.

This is consistent with prior observations that aldosterone regulates urinary kallikrein activity (20). Previous work demonstrated that mice with global deletion of kallikrein-1 exhibited reduced γ-ENaC cleavage and a blunted natriuretic response to amiloride yet maintained overall sodium balance (3). Because this model also displayed vascular and cardiac abnormalities (23, 24), careful interpretation of renal-specific effects is needed. We hypothesized that targeted deletion of kallikrein-1 in the CNT/CCD would unmask a more pronounced defect in ENaC-dependent sodium handling. However, at baseline CNT/CCD-specific *Klk1* knockout mice maintained sodium and potassium homeostasis and γ-ENaC cleavage like their control littermates. Under conditions of sodium restriction and potassium loading, however, *Klk1*-deficient mice displayed a significant reduction in cleaved γ-ENaC protein. Despite the biochemical evidence for impaired ENaC activation during stimulation of aldosterone secretion, amiloride-sensitive natriuresis was preserved in CNT/CCD-specific *Klk1* knockout mice. This finding suggests the presence of compensatory (proteolytic or non-proteolytic) mechanisms that sustain ENaC activity in the absence of kallikrein-1. Indeed, multiple serine proteases, including prostasin and other members of the CAP family (channel-activating proteases), have been implicated in ENaC activation (3, 13), and redundancy among these enzymes may buffer against the loss of any single protease.

Unexpectedly, our urine proteomics analysis did not reveal evidence for compensation by alternative urinary proteases. Instead, kallikrein-1 levels remained among the most abundant urinary proteins in CNT-*Klk1*^-/-^ despite ∼85% less abundance than controls. This apparent discrepancy highlights the disproportionate enrichment of kallikrein-1 in the urinary proteome. One implication is that even lower residual levels of kallikrein-1 may be sufficient to sustain ENaC activation. Indeed, we detected low expression of kallikrein-1 in calbindin negative tubules of CNT-*Klk1*^*-/-*^, from which the protein may be secreted by nephron segments that do not express calbindin in sufficient amounts to activate ENaC in the tubule despite the large reduction in whole kidney kallikrein-1 protein detected by western blot. Alternatively, kallikrein-1 may be secreted downstream of the CNT/CCD or within the lower urinary tract, where it would not be expected to play a direct role in ENaC cleavage. Another consideration is that the mouse genome contains a large subfamily of kallikrein-1 related proteins (Klk1b(x)s) for which there are no orthologous genes in humans, and some of these genes may retain proteolytic activity (31). However, the only such kallikrein-1 related protein we were able to detect in urine was Klk1b5, the quantity of which was unchanged compared with controls.

As *Calb1*-Cre is constitutive, gene recombination occurs early in development, which may allow time for the development of compensatory mechanisms for ENaC regulation. In addition to the kidney, *Calb1* is expressed in some neuronal cell types (32). While it is unlikely to influence urinary kallikrein directly, we cannot rule out the contribution of systemic or off-target effects resulting from neuronal *Klk1* recombination.

Recent studies have challenged a major role for γ-ENaC cleavage for ENaC function *in vivo*. Ray and colleagues demonstrated that mice harboring mutations at the furin cleavage site of γ-ENaC exhibit normal sodium transport and ENaC activity in cortical collecting ducts, despite the absence of detectable cleavage fragments (33). These findings suggest that cleavage-dependent activation of γ-ENaC, while robust *in vitro*, may be dispensable or compensated for *in vivo*. Our own findings align with this conclusion: although γ-ENaC cleavage is clearly diminished in CNT-specific *Klk1* knockout mice, overall sodium and potassium homeostasis is largely preserved, and ENaC function is sufficient to prevent overt sodium wasting. These observations point to a more nuanced role for γ-ENaC cleavage—one that may modulate, rather than act as a sole determinant of, ENaC activity under physiological conditions.

A surprising finding of this study is the observation that deletion of kallikrein-1 led to reduced cleavage of α-ENaC, perhaps suggesting a broader role for kallikrein-1 in modulating ENaC subunit processing. To our knowledge, previous studies of the global *Klk1* knockout did not report on the expression of cleaved α-ENaC species (3). The present study used a well validated α-ENaC antibody from Loffing group ((27), as commercial α-ENaC antibodies may lack specificity (26). Given the lack of a known extracellular cleavage site for α-ENaC by kallikrein-1, this finding is unlikely to be a direct effect of kallikrein-1. Whether the phenomenon is mediated by secondary pathways such as bradykinin remains to be determined. A key function of kallikrein-1 is its kininogenase activity, which generates bradykinin by cleaving low molecular weight kininogen. Bradykinin exerts biological effects primarily through the bradykinin B2 receptor (B2R), which is also expressed in the distal nephron. This raises the possibility that kallikrein-1 may regulate ENaC both directly—through proteolytic cleavage—and indirectly, via bradykinin generation and B2R signaling. Mice globally lacking the B2 receptor exhibit increased γ-ENaC cleavage, but we cannot rule out that this effect is secondary to systemic loss of the receptor rather than loss of B2R in the distal nephron (3). If this effect originated from renal B2R, we would predict that kallikrein-1 deficiency would lead to *increased* ENaC activity due to reduced bradykinin generation. Instead, we observed reduced γ-ENaC cleavage in distal nephron specific *Klk1* knockout mice, and future studies using conditional kidney-specific B2R deletion are needed to fully elucidate the contribution of each of these mechanisms to ENaC regulation.

Taken together, our findings support a model in which aldosterone directs ENaC biology in part by inducing the expression and secretion of kallikrein-1. This protease then facilitates the cleavage of γ-ENaC. However, the physiological significance of this pathway is modulated by redundancy among regulatory mechanisms. Considering emerging data questioning the necessity of ENaC cleavage for function, future studies should focus on dissecting the relative contributions of proteolytic and non-proteolytic mechanisms of ENaC regulation, including the interplay between kallikrein-1, B2R, and the aldosterone axis.

## Supplemental Material

https://doi.org/10.6084/m9.figshare.30166810.v1

## Acknowledgments

We thank Pat Thaitongsuk and Jessica Bahena-Lopez for assistance with collection of samples. The α-ENaC antibody was generously provided by J. Loffing (Institute of Anatomy, University of Zurich, Zurich, Switzerland).

## References

1. Kleyman TR, Kashlan OB, Hughey RP. Epithelial Na(+) Channel Regulation by Extracellular and Intracellular Factors. Annu Rev Physiol. 2018;80:263–81.

2. Frindt G, McNair T, Dahlmann A, Jacobs-Palmer E, Palmer LG. Epithelial Na channels and short-term renal response to salt deprivation. Am J Physiol Renal Physiol. 2002;283(4):F717–26.

3. Picard N, Eladari D, El Moghrabi S, Planes C, Bourgeois S, Houillier P, et al. Defective ENaC processing and function in tissue kallikrein-deficient mice. J Biol Chem. 2008;283(8):4602–11.

4. Nesterov V, Bertog M, Canonica J, Hummler E, Coleman R, Welling PA, et al. Critical role of the mineralocorticoid receptor in aldosterone-dependent and aldosterone-independent regulation of ENaC in the distal nephron. Am J Physiol Renal Physiol. 2021;321(3):F257–F68.

5. Terker AS, Yarbrough B, Ferdaus MZ, Lazelle RA, Erspamer KJ, Meermeier NP, et al. Direct and Indirect Mineralocorticoid Effects Determine Distal Salt Transport. J Am Soc Nephrol. 2016;27(8):2436–45.

6. Maeoka Y, Su XT, Wang WH, Duan XP, Sharma A, Li N, et al. Mineralocorticoid Receptor Antagonists Cause Natriuresis in the Absence of Aldosterone. Hypertension. 2022;79(7):1423–34.

7. Hughey RP, Bruns JB, Kinlough CL, Harkleroad KL, Tong Q, Carattino MD, et al. Epithelial sodium channels are activated by furin-dependent proteolysis. J Biol Chem. 2004;279(18):18111–4.

8. Sheng S, Carattino MD, Bruns JB, Hughey RP, Kleyman TR. Furin cleavage activates the epithelial Na+ channel by relieving Na+ self-inhibition. Am J Physiol Renal Physiol. 2006;290(6):F1488–96.

9. Masilamani S, Kim GH, Mitchell C, Wade JB, Knepper MA. Aldosterone-mediated regulation of ENaC alpha, beta, and gamma subunit proteins in rat kidney. J Clin Invest. 1999;104(7):R19–23.

10. Frindt G, Masilamani S, Knepper MA, Palmer LG. Activation of epithelial Na channels during short-term Na deprivation. Am J Physiol Renal Physiol. 2001;280(1):F112–8.

11. Noreng S, Bharadwaj A, Posert R, Yoshioka C, Baconguis I. Structure of the human epithelial sodium channel by cryo-electron microscopy. Elife. 2018;7.

12. Carattino MD, Sheng S, Bruns JB, Pilewski JM, Hughey RP, Kleyman TR. The epithelial Na+ channel is inhibited by a peptide derived from proteolytic processing of its alpha subunit. J Biol Chem. 2006;281(27):18901–7.

13. Bruns JB, Carattino MD, Sheng S, Maarouf AB, Weisz OA, Pilewski JM, et al. Epithelial Na+ channels are fully activated by furin- and prostasin-dependent release of an inhibitory peptide from the gamma-subunit. J Biol Chem. 2007;282(9):6153–60.

14. Wang XP, Balchak DM, Gentilcore C, Clark NL, Kashlan OB. Activation by cleavage of the epithelial Na(+) channel alpha and gamma subunits independently coevolved with the vertebrate terrestrial migration. Elife. 2022;11.

15. Caldwell RA, Boucher RC, Stutts MJ. Serine protease activation of near-silent epithelial Na+ channels. Am J Physiol Cell Physiol. 2004;286(1):C190–4.

16. Diakov A, Bera K, Mokrushina M, Krueger B, Korbmacher C. Cleavage in the gamma-subunit of the epithelial sodium channel (ENaC) plays an important role in the proteolytic activation of near-silent channels. J Physiol. 2008;586(19):4587–608.

17. Patel AB, Chao J, Palmer LG. Tissue kallikrein activation of the epithelial Na channel. Am J Physiol Renal Physiol. 2012;303(4):F540–50.

18. Vuagniaux G, Vallet V, Jaeger NF, Pfister C, Bens M, Farman N, et al. Activation of the amiloride-sensitive epithelial sodium channel by the serine protease mCAP1 expressed in a mouse cortical collecting duct cell line. J Am Soc Nephrol. 2000;11(5):828–34.

19. Vallet V, Chraibi A, Gaeggeler HP, Horisberger JD, Rossier BC. An epithelial serine protease activates the amiloride-sensitive sodium channel. Nature. 1997;389(6651):607–10.

20. Margolius HS, Horwitz D, Geller RG, Alexander RW, Gill JR, Jr., Pisano JJ, et al. Urinary kallikrein excretion in normal man. Relationships to sodium intake and sodium-retaining steroids. Circ Res. 1974;35(6):812–9.

21. Rhaleb NE, Yang XP, Carretero OA. The kallikrein-kinin system as a regulator of cardiovascular and renal function. Compr Physiol. 2011;1(2):971–93.

22. Rohrwasser A, Ishigami T, Gociman B, Lantelme P, Morgan T, Cheng T, et al. Renin and kallikrein in connecting tubule of mouse. Kidney Int. 2003;64(6):2155–62.

23. Bergaya S, Meneton P, Bloch-Faure M, Mathieu E, Alhenc-Gelas F, Levy BI, et al. Decreased flow-dependent dilation in carotid arteries of tissue kallikrein-knockout mice. Circ Res. 2001;88(6):593–9.

24. Meneton P, Bloch-Faure M, Hagege AA, Ruetten H, Huang W, Bergaya S, et al. Cardiovascular abnormalities with normal blood pressure in tissue kallikrein-deficient mice. Proc Natl Acad Sci U S A. 2001;98(5):2634–9.

25. El Moghrabi S, Houillier P, Picard N, Sohet F, Wootla B, Bloch-Faure M, et al. Tissue kallikrein permits early renal adaptation to potassium load. Proc Natl Acad Sci U S A. 2010;107(30):13526–31.

26. Mutchler SM, Shi S, Whelan SCM, Kleyman TR. Validation of commercially available antibodies directed against subunits of the epithelial Na(+) channel. Physiol Rep. 2023;11(1):e15554.

27. Sorensen MV, Grossmann S, Roesinger M, Gresko N, Todkar AP, Barmettler G, et al. Rapid dephosphorylation of the renal sodium chloride cotransporter in response to oral potassium intake in mice. Kidney Int. 2013;83(5):811–24.

28. Lee CT, Ng HY, Lee YT, Lai LW, Lien YH. The role of calbindin-D28k on renal calcium and magnesium handling during treatment with loop and thiazide diuretics. Am J Physiol Renal Physiol. 2016;310(3):F230–6.

29. Bostanjoglo M, Reeves WB, Reilly RF, Velazquez H, Robertson N, Litwack G, et al. 11Beta-hydroxysteroid dehydrogenase, mineralocorticoid receptor, and thiazide-sensitive Na-Cl cotransporter expression by distal tubules. J Am Soc Nephrol. 1998;9(8):1347–58.

30. McCormick JA, Mutig K, Nelson JH, Saritas T, Hoorn EJ, Yang CL, et al. A SPAK isoform switch modulates renal salt transport and blood pressure. Cell Metab. 2011;14(3):352–64.

31. Moustardas P, Yamada-Fowler N, Apostolou E, Tzioufas AG, Turkina MV, Spyrou G. Deregulation of the Kallikrein Protease Family in the Salivary Glands of the Sjogren’s Syndrome ERdj5 Knockout Mouse Model. Front Immunol. 2021;12:693911.

32. Zhang B, Li L, Tang X, Zeng J, Song Y, Hou Z, et al. Distribution Patterns of Subgroups of Inhibitory Neurons Divided by Calbindin 1. Mol Neurobiol. 2023;60(12):7285–96.

33. Ray EC, Nickerson A, Sheng S, Carrisoza-Gaytan R, Lam T, Marciszyn A, et al. Influence of proteolytic cleavage of ENaC’s gamma subunit upon Na(+) and K(+) handling. Am J Physiol Renal Physiol. 2024;326(6):F1066–F77.

