## Supplementary Materials for "Kidney kallikrein-1 contributes to cleavage of gamma-ENaC *in vivo*"

**Supplemental Table 1. List of antibodies used in experiments**

| **Target** | **Host Species** | **Dilution (WB)** | **Dilution (IF)** | **Source** | **Reference** |
| --- | --- | --- | --- | --- | --- |
| γ-ENaC | Rabbit | 1:1000 | 1:500 | Stressmarq #SPC-405D | (1) |
| α-ENaC | Rabbit | 1:5000 |  | Loffing lab | (2) |
| Kallikrein-1 | Rabbit | 1:1000 | 1:1000 | Boster #PA1709 | This study |
| Total NCC | Rabbit | 1:2000 |  | Ellison laboratory | (3) |
| pNCC T53 | Rabbit | 1:2000 |  | Ellison laboratory | (4) |
| Calbindin-D28k | Mouse |  | 1:500 | Swant | (5) |

**Supplementary Table 2. Physiological parameters from controls and connecting tubule specific kallikrein-1 knockout mice housed in metabolic cages.**

|  | **CNT *Klk1*^+/+^** | | | **CNT *Klk1*^-/-^** | | | **P Value** |
| --- | --- | --- | --- | --- | --- | --- | --- |
|  | **Mean** | **SEM** | ***n*** | **Mean** | **SEM** | ***n*** |  |
| **Male** | | | | | | | |
| ***Control Diet*** |  |  |  |  |  |  |  |
| Age (wk) | 13.1 | 0.67 | 8 | 12.4 | 1.44 | 5 | 0.61 |
| Weight (g) | 27.8 | 0.35 | 8 | 25.8 | 0.59 | 5 | 0.06 |
| Food (g) | 8.08 | 1.59 | 8 | 5.81 | 1.39 | 5 | 0.35 |
| Volume (mL/24hr) | 2.18 | 0.19 | 8 | 2.09 | 0.23 | 5 | 0.77 |
| ***Low Na/High K Diet*** |  |  |  |  |  |  |  |
| Age (wk) | 12.7 | 1.32 | 7 | 12.0 | 1.05 | 7 | 0.68 |
| Weight (g) | 25.8 | 0.59 | 7 | 25.5 | 0.95 | 7 | 0.79 |
| Food (g) | 9.50 | 1.74 | 7 | 6.80 | 1.43 | 7 | 0.25 |
| Volume (mL/24hr) | 2.69 | 0.23 | 7 | 2.71 | 0.33 | 7 | 0.97 |
| **Female** | | | | | | | |
| ***Control Diet*** |  |  |  |  |  |  |  |
| Age (wk) | 14.6 | 0.40 | 5 | 13.0 | 1.30 | 5 | 0.27 |
| Weight (g) | 20.3 | 0.86 | 5 | 19.7 | 1.17 | 5 | 0.72 |
| Food (g) | 9.67 | 1.42 | 5 | 7.31 | 0.92 | 5 | 0.20 |
| Volume (mL/24hr) | 2.48 | 0.31 | 5 | 2.57 | 0.36 | 5 | 0.86 |
| ***Low Na/High K Diet*** |  |  |  |  |  |  |  |
| Age (wk) | 11.0 | 1.35 | 4 | 12.0 | 1.41 | 6 | 0.64 |
| Weight (g) | 20.0 | 0.87 | 4 | 20.8 | 0.70 | 6 | 0.55 |
| Food (g) | 5.09 | 1.38 | 4 | 5.66 | 1.74 | 6 | 0.81 |
| Volume (mL/24hr) | 2.42 | 0.10 | 4 | 1.80 | 0.37 | 6 | 0.23 |

**Supplementary Table 3. Blood parameters of female mice on control and low sodium / high potassium diets**

|  | **CNT *Klk1*^+/+^** | | | **CNT *Klk1*^-/-^** | | | **P Value** |
| --- | --- | --- | --- | --- | --- | --- | --- |
|  | **Mean** | **SEM** | ***n*** | **Mean** | **SEM** | ***n*** |  |
| **Parameter** | | | | | | | |
| ***Control Diet*** | |  |  |  |  |  |  |
| Na | 139.0 | 3.00 | 5 | 144.0 | 1.30 | 5 | 0.14 |
| K | 4.12 | 0.37 | 5 | 3.80 | 0.05 | 5 | 0.36 |
| Cl | 113.8 | 3.09 | 5 | 115.8 | 2.78 | 5 | 0.64 |
| TCO2 | 16.75 | 1.18 | 5 | 19.0 | 1.18 | 5 | 0.23 |
| iCa | 1.19 | 0.04 | 5 | 1.28 | 0.01 | 5 | 0.03 |
| Glu | 287.8 | 15.93 | 5 | 226.0 | 30.96 | 5 | 0.15 |
| BUN | 25.00 | 2.35 | 5 | 26.20 | 3.22 | 5 | 0.78 |
| Hct | 38.00 | 0.71 | 5 | 36.80 | 0.92 | 5 | 0.35 |
| ***Low Na/High K Diet*** | |  |  |  |  |  |  |
| Na | 142.8 | 1.11 | 4 | 141.8 | 0.95 | 6 | 0.55 |
| K | 4.80 | 0.43 | 4 | 3.97 | 0.21 | 6 | 0.09 |
| Cl | 114.8 | 2.14 | 4 | 113.0 | 1.37 | 6 | 0.49 |
| TCO2 | 18.00 | 1.08 | 4 | 18.17 | 0.75 | 6 | 0.90 |
| iCa | 1.31 | 0.03 | 4 | 1.28 | 0.00 | 6 | 0.16 |
| Glu | 256.00 | 20.10 | 4 | 326.67 | 12.05 | 6 | 0.01 |
| BUN | 22.25 | 1.38 | 4 | 19.67 | 1.54 | 6 | 0.28 |
| Hct | 37.50 | 1.04 | 4 | 35.67 | 0.61 | 6 | 0.14 |

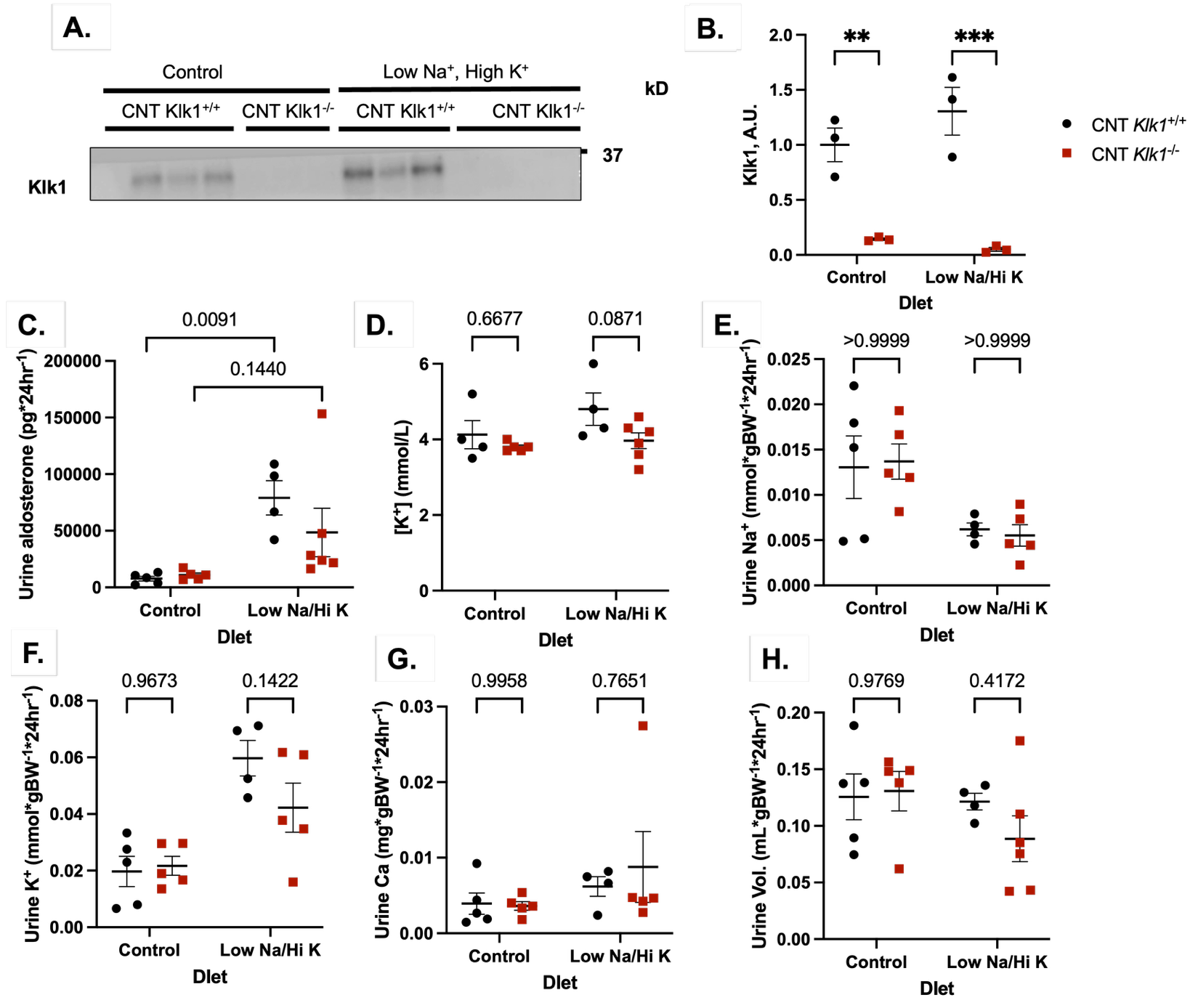

**Supplemental Figure 1. Female Mice with Distal Nephron-Specific Deletion of Kallikrein-1 Maximally Conserve Potassium in response to a Low Sodium, High Potassium Diet.** These results recapitulate our findings in male mice (Fig 1). A. Western blot kallikrein-1 demonstrating a significant reduction in kallikrein-1 expression in the *Klk1*^flox/flox^ *Calb1*-Cre mice (CNT *Klk1*^-/-^) follow 5 days on a normal salt diet (Control) or low sodium, high potassium diet (Low Na/Hi K). On the final day of dietary challenge, mice were placed in metabolic cages for 24 hour urine collection then harvested for blood and kidney tissue. C. 24-hour urinary aldosterone excretion. D. Serum potassium was trending lower in CNT *Klk1*^-/-^ on the Low Na/Hi K diet. We were unable to detect a difference in urine volume (H) or urinary sodium (E), potassium (F), or calcium (G) excretion under either dietary condition. Results were analyzed by two-way ANOVA followed by Bonferroni multiple comparison correction, with numerical *P* values shown in brackets above each comparison.

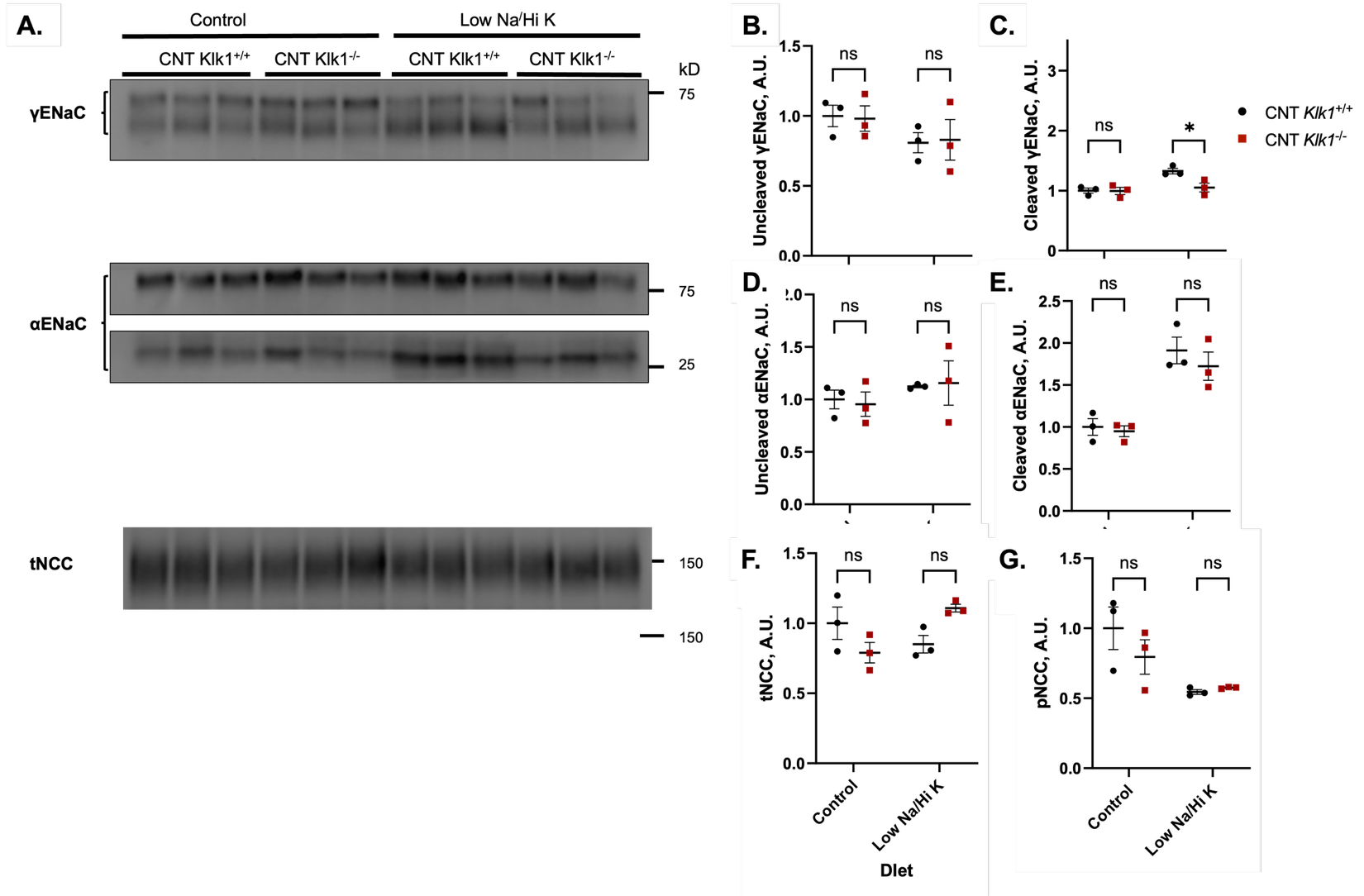

**Supplemental Figure 2. Females largely recapitulate our findings in male CNT *Klk1^-/-^* mice.**

A. Western blot of γ-ENaC, α-ENaC, total NCC (tNCC) and phosphorylated NCC (pNCC) abundance. B-G. Quantification of western blot results. A significant reduction in cleaved γ-ENaC was detected in CNT-*Klk1*^-/-^ mice following challenge with a low sodium, high potassium diet (Low Na/Hi K). Results were analyzed by two-way ANOVA followed by Bonferroni multiple comparison correction. **P*<0.05.

**References**

1.	Mutchler SM, Shi S, Whelan SCM, Kleyman TR. Validation of commercially available antibodies directed against subunits of the epithelial Na(+) channel. Physiol Rep. 2023;11(1):e15554.

2.	Sorensen MV, Grossmann S, Roesinger M, Gresko N, Todkar AP, Barmettler G, et al. Rapid dephosphorylation of the renal sodium chloride cotransporter in response to oral potassium intake in mice. Kidney Int. 2013;83(5):811-24.

3.	Bostanjoglo M, Reeves WB, Reilly RF, Velazquez H, Robertson N, Litwack G, et al. 11Beta-hydroxysteroid dehydrogenase, mineralocorticoid receptor, and thiazide-sensitive Na-Cl cotransporter expression by distal tubules. J Am Soc Nephrol. 1998;9(8):1347-58.

4.	McCormick JA, Mutig K, Nelson JH, Saritas T, Hoorn EJ, Yang CL, et al. A SPAK isoform switch modulates renal salt transport and blood pressure. Cell Metab. 2011;14(3):352-64.

5.	Lee CT, Ng HY, Lee YT, Lai LW, Lien YH. The role of calbindin-D28k on renal calcium and magnesium handling during treatment with loop and thiazide diuretics. Am J Physiol Renal Physiol. 2016;310(3):F230-6.
